# Red2Flpe-SCON: A Versatile, Multicolor Strategy for Generating Mosaic Conditional Knockout Mice

**DOI:** 10.1101/2023.02.09.527641

**Authors:** Szu-Hsien Sam Wu, Somi Kim, Heetak Lee, Ji-Hyun Lee, Gabriele Colozza, So-Yeon Park, Réka Bakonyi, Isaree Teriyapirom, Natalia Hallay, Sandra Pilat-Carrota, Hans-Christian Theussl, Jihoon Kim, Joo-Hyeon Lee, Benjamin D. Simons, Jong Kyoung Kim, Bon-Kyoung Koo

## Abstract

Image-based lineage tracing enables tissue turnover kinetics and lineage potentials of different adult cell populations to be investigated. Previously, we reported a genetic mouse model system, *Red2Onco*, which ectopically expressed mutated oncogenes together with red fluorescent proteins (RFP). This system enabled the expansion kinetics and neighboring effects of oncogenic clones to be dissected. We now report Red2Flpe-SCON: a new mosaic knockout system that uses multicolor reporters to label both mutant and wild-type cells. We have developed the *Red2Flpe* mouse line for red clone-specific Flpe expression, as well as the FRT-based SCON (<u>S</u>hort <u>C</u>onditional Intr<u>ON</u>) method to facilitate tunable conditional mosaic knockouts in mice. We used the Red2Flpe-SCON method to study Sox2 mutant clonal analysis in the esophageal epithelium of adult mice which revealed that the stem cell gene, Sox2, is not essential for adult stem cell maintenance itself, but rather for stem cell proliferation and differentiation.

## Introduction

Adult tissue homeostasis is regulated by resident stem cells, which divide at various frequencies to either self-renew or differentiate. Competition between stem cells for niche spaces occurs in many organs throughout life (Martincorena et al. 2015, 2018; Lee-Six et al. 2018). With advances in sequencing technologies and model systems, it has become clear that cell competition plays an important role in regulating tissue turnover and cancer development (Colom et al. 2021). While prospective or retrospective computational-based lineage reconstruction enables high-throughput analyses of clonal information from tissues, the precise spatial distribution and morphology of the clones can only be revealed using image-based lineage tracing (Wu et al. 2019).

Multicolor reporter systems have been crucial to understanding the turnover kinetics that occur during tissue homeostasis (Snippert et al. 2010; Lopez-Garcia et al. 2010; Doupé et al. 2012; Han et al. 2019). Combining fluorescent reporters with conditional gene knockout (cKO) alleles enables gene knockouts and lineage tracing to be carried out simultaneously (Vermeulen et al. 2013; Snippert et al. 2014), while utilizing two recombinases in a single mouse provides further control (Lao et al. 2012; Thorsen et al. 2021). However, such an approach carries the risk of genotype-phenotype mismatches. Although approaches that use in utero or in vivo electroporation (Kim et al. 2019) and transduction (Kohara et al. 2020) have been developed — circumventing the need for new mouse lines — they are fundamentally limited, as they target cell populations in an unspecific manner.

To achieve precise mosaic genetic labeling and tracing, several systems have been developed including, mosaic analysis with double markers (MADM) (Zong et al. 2005; Contreras et al. 2021), IfgMosaics (Pontes-Quero et al. 2017) and Red2Onco (Yum et al. 2021). These systems provide a near-definitive genotype-phenotype correlation. Moreover, in these mosaic systems, both mutant and wild-type clones are fluorescently labeled; this enables direct comparisons between lineage kinetics and any potential non-cell-autonomous effects (Table 1). Although mosaic knockouts are achievable using MADM, this method relies on the inefficient inter-chromosomal recombination events that require cell division (Zong et al. 2005). The MADM and Red2cDNA systems both have reproducible initial labeling ratios between the different colors; however, quantitative analysis is not possible with the ifgMosaics system, as it yields variable labeling ratios (Table 1).

**Table 1.**
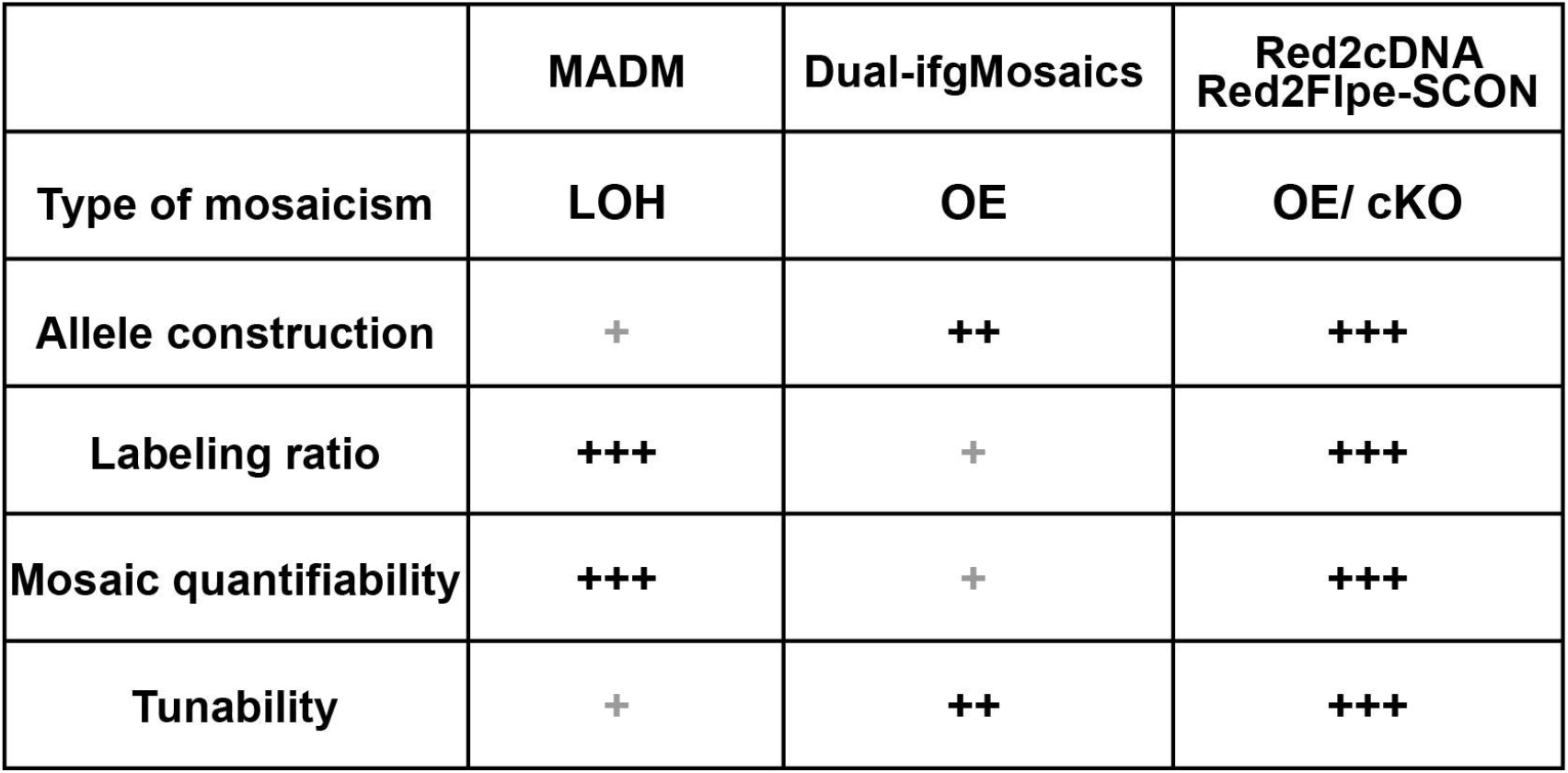
Comparison of different in vivo mosaic genetic systems.

While mosaic overexpression systems are useful for studying specific decisions regarding cell fate (Pontes-Quero et al. 2017) and/or oncogenic activations (Yum et al. 2021), a mosaic knockout approach is more suitable for studying general effects on the system in various biological contexts. In this study, we report a new, tunable and efficient mosaic knockout system: Red2Flpe-SCON. This system is based on the Confetti reporter system and uses red clone-specific Flpe expression with FRT-based Short Conditional Intr<u>ON</u> (SCON). Our system offers a versatile, tunable, and precise method to achieve Flp-based mosaic knockout with multicolor labeling in mice.

## Results

### Generating Red2cDNA: a mosaic system based on the Confetti reporter allele using CRISPR nickase-mediated targeting

We previously reported the *Red2Onco system —* a series of modified Rosa26-Confetti (Brainbow2.1) alleles, which enable the mosaic, ectopic expression of oncogenes in a red fluorescent protein (RFP)-labeled, clone-specific manner (Yum et al. 2021). We adapted an efficient electroporation protocol to achieve the desired gene knock-in, adjacent to the RFP, using CRISPR-Cas9 nickases with Confetti embryonic stem cells (ESCs); a process that usually takes approximately two to three weeks (Figure S1). The targeting approach harnesses the homology-directed repair (HDR) pathway, with homology arms of 600-700bp in length. The Red2 targeting vector contains specific restriction cloning sites for inserting the cDNA sequence — downstream and in frame with the RFP and 2A peptide — and a PGK-Blasticidin-pA cassette with an inverted orientation for efficient antibiotic selection (Figure S1A). The RFP protein, *tdimer2*, of the confetti allele consists of two segments with repeated sequences. The left homology arm contains the sequence of one dimeric unit and can therefore be inserted to replace either dimer (with about 650bp difference in length), where the second dimer is the desirable target (Figure S1) (Campbell et al. 2002). For the targeting experiment, we utilized a pair of sgRNAs with a Cas9-D10A nickase (which only cleaves strands that are complementary to the sgRNA) to minimize any potential off-target effects (Figure S1C) (Ran et al. 2013). Upon electroporation, a small fraction of cells showed GFP expression from the Cas9 nickase vectors (Figure S1D). After 48-72hr of recovery, cells were subjected to blasticidin treatment to select for targeted clones. The verified clones were checked by long range PCR genotyping (Figure S1E) and were then subsequently injected into developing blastocysts to generate chimeras (Figure S1F). This CRISPR-mediated method of targeting enabled mosaic genetic Red2cDNA mouse lines to be generated efficiently.

### Red2-Flpe: a versatile tool that enables precise mosaic knockout with multicolor labeling

Here, we present a mosaic knockout system that involves the insertion of the Flpe recombinase sequence (Buchholz et al. 1998),(Buchholz et al. 1998),(Buchholz et al. 1998), adjacent to an RFP (tdimer2) linked with a P2A peptide. This setup allows for the RFP-labeled (red), clone-specific, Flpe-mediated recombination of the target FRT allele, while ensuring that all the other fluorescent proteins remain genetically unaltered (Figure 1). We named this system, *Red2Flpe*.

**Figure 1.**
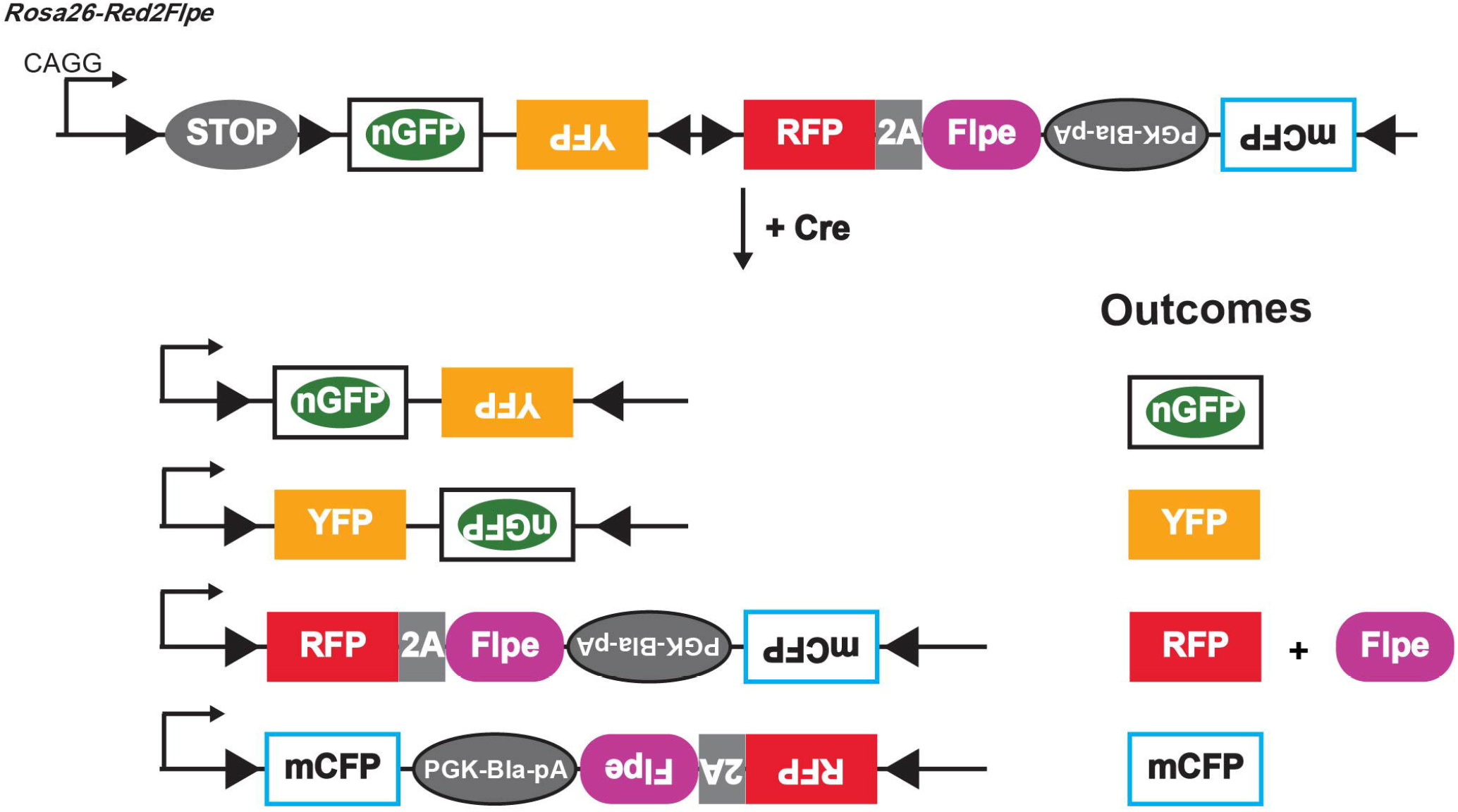
Design of theRed2Flpe system: a mosaic knockout multicolor reporter allele. Upon Cre induction, each cell (unless it is polyploid) is adapted to express one fluorescent protein color. The cells labeled with red fluorescent protein (RFP) express the Flpe recombinase, while all the other fluorescent protein colors (green/ yellow/ cyan) correspond to wildtype cells.

After activating *Red2Flpe* with the ubiquitous *Rosa-CreER^T2^* line, we harvested the seminal vesicles (Figure S2A–S2D), spleen (Figure S2E–S2H) and tongue (Figure S2I–S2L) and confirmed that YFP and RFP signals were present. This demonstrated the utility of *Red2Flpe* in multiple organs and tissues with recombination rates that were suitable for mosaic lineage tracing studies.

We next tested whether the expression of Flpe on the modified Confetti allele could recombine with the FRT sites on different alleles in the mouse genome, in a red clone-specific manner. In combination with the intestinal epithelium-specific CreER^T2^ line, *Villin-CreERT^2^*, we crossed *Red2Flpe* with two separate FRT lines. Firstly, *Vil-CreER^T2^;Red2Flpe* was crossed with a newly generated Apc-FRT (exon 15 flanked by FRT sites) (Figure 2A); knockout of this leads to stabilized β-catenin accumulation and subsequent Wnt signaling activation (Colnot et al. 2004). One week after the Cre-mediated labeling (Figure 2B), the *Vil-CreER^T2^;Red2Flp;Apc^FRT/FRT^* mouse intestine showed that the RFP clones, specifically, displayed elevated cytoplasmic β-catenin staining (Figure 2C). Secondly, the *Vil-CreER^T2^;Red2Flpe* line was crossed with an FRT-based GFP reporter line called *RCE:FRT*(an FRT-stop-FRT GFP reporter on the Rosa26 locus) (Figure 2D) (Sousa et al. 2009). We expected overlapping GFP and RFP signals but no overlap with any other colors, if Flpe worked efficiently and specifically. After one week of Cre induction, we only observed GFP expression in the Flpe-expressing RFP+ cells (Figure 2E and 2F). To assess the efficiency and specificity of Flpe/FRT recombination of Red2Flpe, primary intestinal cells were harvested and analyzed by flow cytometry at five, seven and ten days after tamoxifen injection (Figure 2G and 2H). A stringent gate was set for the RFP-positive cells, as the maturation time is longer for tdimer2 (~120min) (Campbell et al. 2002) (Figure 2G). We observed an increase in the proportion of GFP/RFP double-positive cells from day five (66.3%) to day seven (82.1%) and day ten (82.6%) (Figure 4H). There is a time lag between the point of tamoxifen injection, the observed Cre activity and the subsequent stable expression of the fluorescent reporters and Flpe recombinase; this data, therefore, suggests that Red2Flpe becomes active approximately five to seven days after the tamoxifen injection (Figure 2H). Overall, these results indicate that *Red2Flpe* is a functional and efficient genetic tool that can be used to generate mosaic multicolor mouse models in vivo.

**Figure 2.**
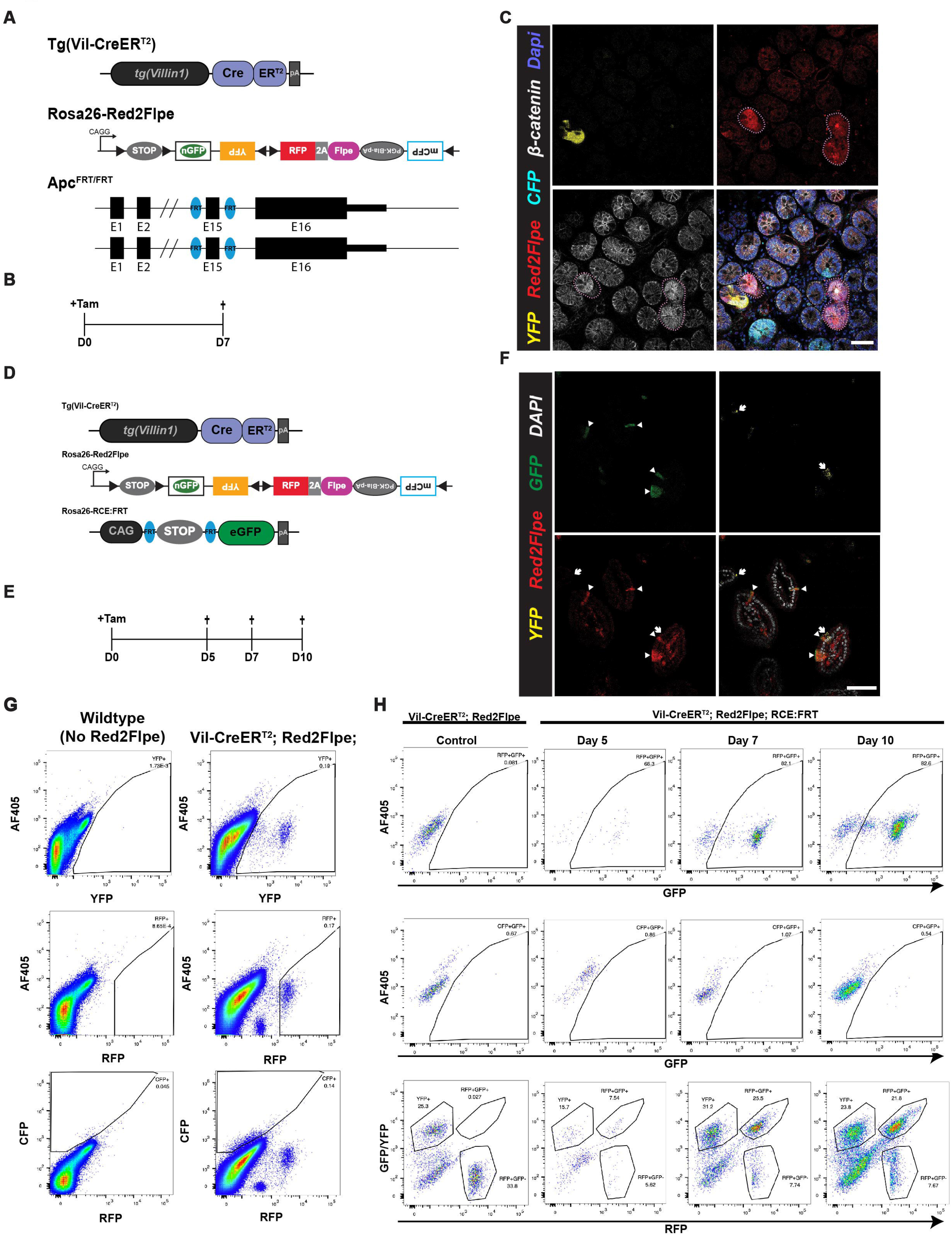
Red2Flpe enables efficient red clone-specific recombination in vivo. **A-C,** Small intestine samples harvested one week after treatment with tamoxifen (2 mg tamoxifen per 20 g body weight) from a Tg(Vil-CreER^T2^); Red2Flpe; Apc^FRT/FRT^ mouse. The wild-type crypts labelled with either cyan (CFP), yellow (YFP), or no color show clear β-catenin staining in the membrane with less staining in the cytoplasm. The RFP-labelled (red) crypts display an increase in the cytoplasmic fraction of β-catenin staining — indicating successful knock out of *Apc* and malfunction of the β-catenin destruction complex. The RFP-labelled (red) crypts and the corresponding β-catenin staining are marked with dotted lines. Scale bar, 50 μm. **D-F.** Small intestine samples harvested one week after treatment with tamoxifen (2 mg tamoxifen per 20 g body weight) from a Tg(Vil-CreER^T2^); Red2Flpe; Gt(ROSA)26Sor^tm1.2(CAG-EGFP)Fsh^ mouse. The expression of RFP (red) coincides with GFP (green) but does not coincide with Confetti YFP (yellow). RFP-GFP double positive cells are marked by the triangles and the YFP cells are marked by the arrows. **G.** Gating strategy of YFP, RFP and CFP cells in tamoxifen-induced Tg(Vil-CreER^T2^); Red2Flpe intestine. **H.** GFP+ cells within the gated RFP+ population (top row), GFP+ cells within the gated CFP+ population (middle row), and the overall view of G/YFP and RFP levels in cells labelled with a color (G/YFP, RFP or CFP) (bottom row) in control (Tg(Vil-CreER^T2^); Red2Flpe), or in an FRT-based reporter (Tg(Vil-CreER^T2^); Red2Flpe; Gt(ROSA)26Sor^tm1.2(CAG-EGFP)Fsh^) mouse harvested 5, 7 or 10 days post tamoxifen induction.

### Short Conditional intrON (SCON) facilitate the generation of FRT-based conditional knockout mouse lines with a one-step zygote injection

As Cre-loxP was found to work more efficiently than the wild-type Flp, most of the murine conditional knockout (cKO) lines were made using the Cre-loxP system. New conditional mouse lines are often generated by in vitro ESC targeting, followed by a blastocyst injection to acquire chimeric mice, and final germline transmission; this process takes at least six months. Despite efforts to improve the efficiency of the zygote injections, using CRISPR-based knock-in with a long single-stranded oligonucleotide (ssODN) (Quadros et al. 2017), the process remains technically challenging.

To facilitate cKO mouse generation, we recently developed an artificial intron-based approach that uses a <u>S</u>hort <u>C</u>onditional intrON (SCON), which is just 189 bp in length (Wu et al. 2022). This method only requires a synthesized oligo template, a synthesized gRNA, and a commercially available Cas protein and/or mRNA. All these components are injected into zygotes to generate a cKO. SCON has a neutral effect following the initial insertion of the target gene, and induces the expected loss of function effect upon recombination in vivo. SCON, therefore, offers an alternative but efficient way to generate cKO alleles using a method that is as simple as CRISPR-based ‘tagging’.

We reasoned that the loxP recombination sites used with SCON could be exchanged with FRTs, for compatibility with the Red2Flpe mosaic knockout system. The SCON-FRT system works in the same way as the SCON-loxP system — which consists of a splice donor, branch point, and splice acceptor — with SCON acting as a functional intron. With this system, two FRT recombination sites flank the branch point; the removal of the branch point upon Flp-mediated recombination then abrogates the SCON’s intronic function, causing it to be retained in the mature transcript after splicing. The remaining 55 bp SCON intron sequence contains potential stop codons that can cause premature termination of translation and subsequent truncation of the target protein (Figure 3A).

**Figure 3.**
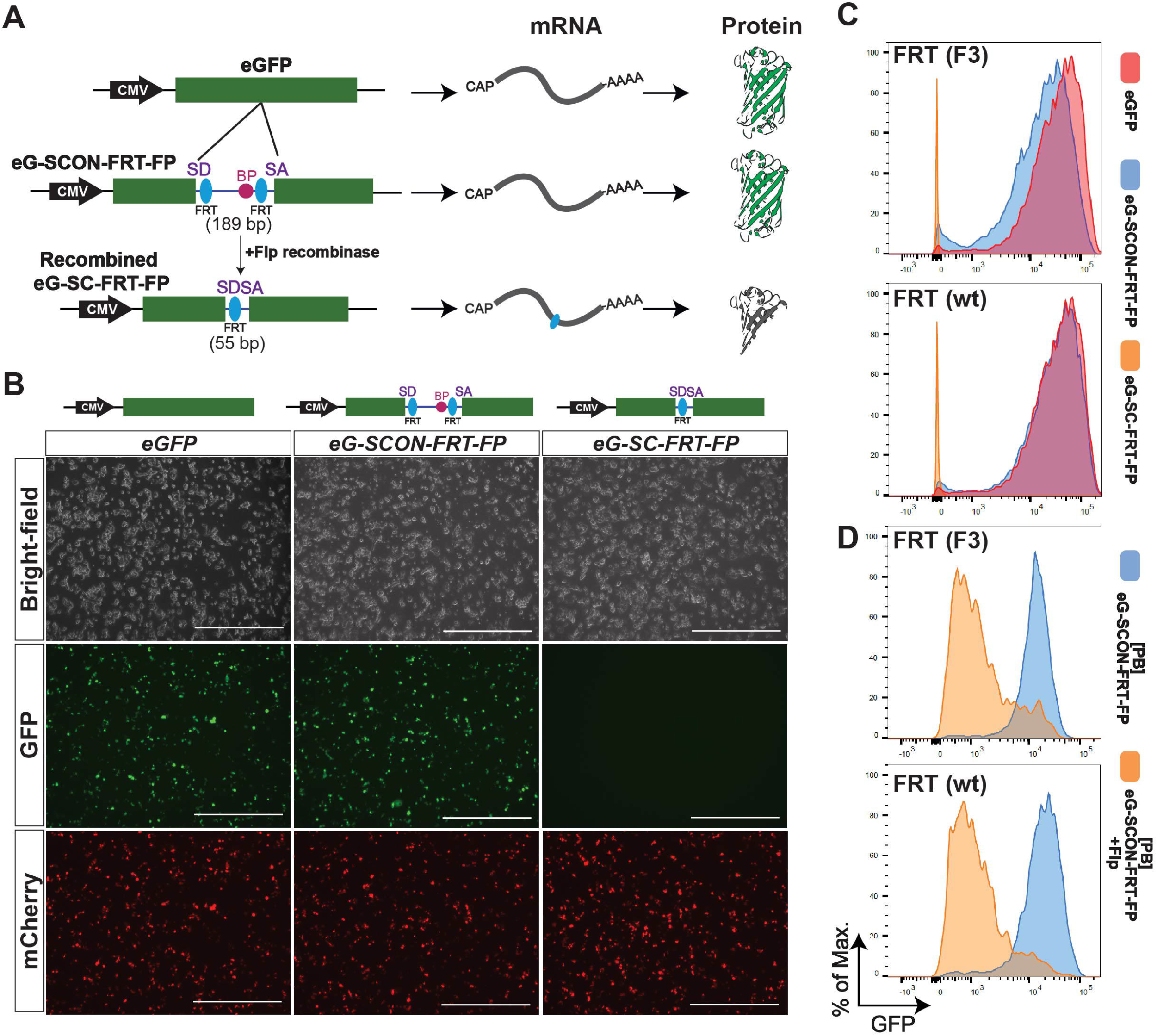
SCON-FRT is a Flpe/FRT-based conditional knockout system. **A.** Schematic diagram of SCON-FRT in an eGFP overexpression construct, including eGFP, eG-SCON-FRT-FP and recombined eG-SCFRT-FP. SD (splice donor), BP (branch point), SA (splice acceptor). **B.** Brightfield and fluorescent images of HEK293T cells 24 hours after transfection. **C.** GFP fluorescence level of HEK293T cells 48 hours after transfection. Red: intact eGFP; Blue: eG-SCON-FRT-FP; Orange: recombined eG-SC-FRT-FP. **D.** GFP fluorescence level of HEK293T cells with integrated eG-SCON-FRT-FP before (blue) and after (orange) transfection of a Flp-expressing plasmid.

We next tested whether the SCON-FRT system had a similar neutral effect on gene expression, as with SCON-loxP. We transfected HEK293T cells with either intact eGFP; eG-SCON-FRT-FP (eGFP with an SCON-FRT insertion); or the recombined form, eG-SC-FRT-FP (Figure 3B). We found that the intact eGFP and the two different eG-SCON-FRT-FPs, with either wildtype or F3 FRT sites, had comparable GFP levels; however, the wildtype FRT version slightly out-performed the F3 FRT version (Figure 3C). With the recombined forms, the level of GFP fluorescence was not detectable, indicating loss of expression (Figure 3B and 3C). We also tested whether SCON-FRT could be efficiently recombined in mammalian cells upon Flp expression. We generated mouse ESCs with constitutive expression of eG-SCON-FRT-FP, using the piggyBac transposon system. After transfecting an Flp-expressing plasmid, we found that the level of GFP diminished significantly, which confirmed the compatibility of the SCON-FRT for Flp/FRT-based recombination system in mammalian cells (Figure 3D). Therefore, we concluded that the SCON-FRT system was also neutral, like the SCON-loxP system, and it was applicable in mammalian cells.

### Red2Flpe can be efficiently utilized in combination with the SCON-FRT system

Next, we assessed the compatibility of the SCON-FRT system with *Red2Flpe*. We made use of *Confetti* and *Red2Flpe* ESCs in combination with piggyBac-eG-SCON-FRT-FPs (Figure 4A). Cells with an integrated eG-SCON-FRT-FP were selected with puromycin with the expectation that both eGFP and puromycin-resistance expression would be compromised following Flp-mediated FRT recombination (Figure 4A and 4B). After selecting GFP-expressing cells, we induced the recombination of the *Confetti* and *Red2Flpe* alleles, respectively, by transfecting a Cre-expressing plasmid. Using fluorescent activated cell sorting (FACS), RFP+ cells were sorted and cultured separately from the uninduced cells. All the RFP+ colonies of *Red2Flpe* ESCs showed no eGFP expression compared to the *Confetti* ESCs, which were used as controls (Figure 4B). Flow cytometry revealed that most of the cells in the uninduced cultures retained high levels of GFP expression, despite some transgene silencing, while all of the recombined RFP+ cells lost GFP expression completely (Figure 4C). These results indicated that Red2Flpe could be coupled with SCON-FRT to successfully achieve conditional mosaic gene knockouts.

**Figure 4.**
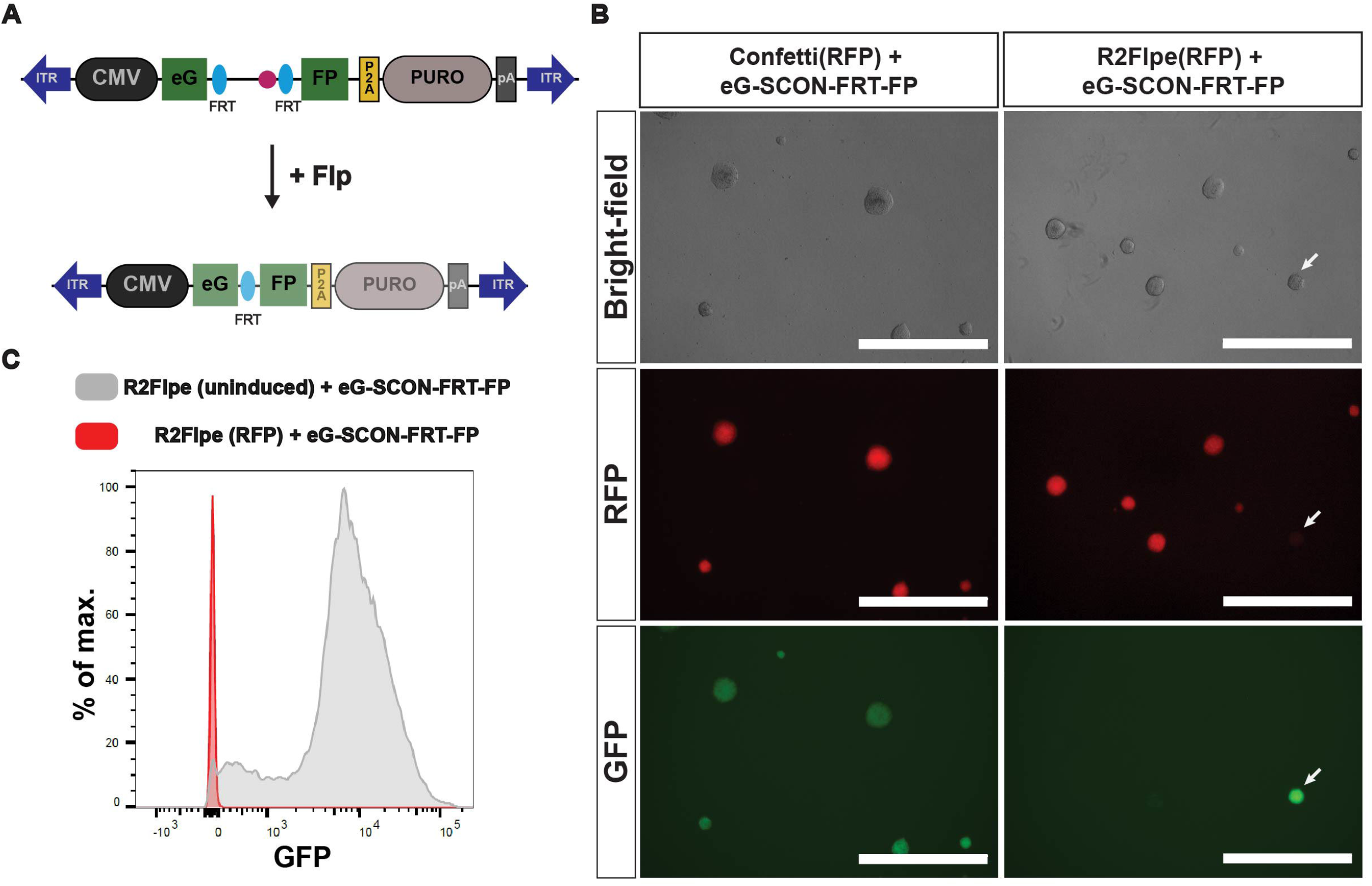
Red2Flpe is compatible with the SCON-FRT system in vitro. **A.** A piggybac vector carrying the overexpression cassette of eG-SCON-FRT-FP coupled with puromycin resistance. Both eGFP and puromycin expression are reduced following Flp/FRT recombination. **B.** Expectations of Red2Flpe; eG-SCON-FRT-FP cells: uninduced or YFP/CFP+ cells retain GFP expression following puromycin withdrawal, and the RFP+ cells that co-express Flpe recombinase lose eGFP expression. **C.** Brightfield and fluorescent images of RFP Confetti and Red2Flpe ESCs with integrated eG-SCON-FRT-FP. The uninduced clone that retains eGFP expression is marked by an arrow. **D**. GFP fluorescence level of Red2Flpe ESCs with integrated eG-SCON-FRT-FP after puromycin withdrawal. Grey: uninduced Red2Flpe; eG-SCON-FRT-FP cells; Red: RFP+ cells.

### Sox2-SCON-FRT mouse generation via one-step zygote injection

With the knowledge that the use of SCONs would not affect basal gene expression, we generated thirteen cKO mouse lines (Wu et al. 2022). We injected CRISPR-Cas9 ribonucleoprotein (RNP), Cas9 mRNA, and a 300 bp long ssODN of SCON (using either loxP sites or FRT sites, both of which were 189 bp long). We used left and right homology arms that were 55 and 56 bp long, respectively (Wu et al. 2022). With the SCON approach, a complete experimental mouse line for mosaic knockout studies could be generated quickly and easily.

We propose a pipeline for generating zygotes using SCON targeting from the desired *CreER* and *Red2Flpe* lines, both in homozygosity (Figure S3A). From the offspring, pups that are heterozygous for the SCON-FRT knock-in can be used for mating to create further experimental lines (Figure S3B). The chance of acquiring the desired experimental cohort is 1/4 (Figure S3C) so, theoretically, it is possible to generate SCON-based *Red2Flpe* mosaic knockout mice by zygote injection with just two mouse generations.

One of the first SCON-FRT lines we generated was the *Sox2-SCON-FRT (Sox2^scon^*) line (Figure S4A). From twenty founder pups, we obtained three heterozygous SCON-FRT knock-in mice with precise integration (15% efficiency). *Sox2^+/scon^* mice can be bred to be homozygous without any noticeable developmental defects (refer to Fig. 4g and h in Wu et al. 2022), which validates the utility of SCON-FRT for in vivo experiments (Wu et al. 2022). The mouse ESCs derived from homozygous *Sox2^scon/scon^* blastocysts expanded stably in culture. Then, following transient Flpe expression, the colonies that contained either mosaic or complete Sox2-KO cells exhibited distorted morphologies. These results were consistent with the importance of Sox2 in maintaining pluripotency (Figure S4B) (Masui et al. 2007).

### Mosaic knockout of Sox2 in adult tissues reveals its variable essentiality

*Sox2* is a transcription factor that plays crucial roles during embryonic development — from the blastocyst stages to the fate-specifying stages — of many tissues (Sarkar and Hochedlinger 2013). Knockout studies revealed that *Sox2* is required for proper development of the esophagus (Que et al. 2007; Trisno et al. 2018). In adults, *Sox2* is expressed in the esophagus and stomach, and is thought to be important for stem cell maintenance (Arnold et al. 2011; DeWard et al. 2014). Consistently, tissue-wide knockout of Sox2 or depletion of Sox2-expressing cells leads to compromised tissue maintenance and physiology (Arnold et al. 2011; DeWard et al. 2014). However, as widespread knockout of Sox2 in these tissues compromises their overall integrity, the exact function of Sox2 remains unclear. A mosaic analysis of Sox2 in the adult esophagus and other tissues is therefore necessary.

We used *Red2Flpe* and the new *Sox2^scon^* alleles with the ubiquitous *Rosa26-CreER^T2^* inducer line, and generated *Rosa26-CreER^T2^; Red2Flpe*; *Sox2^scon/scon^* (*Red2Sox2KO*) mice to investigate the function of Sox2 in the adult mouse esophagus (Figure 5A). By administering tamoxifen to the *Red2Sox2KO* mice, we activated fluorescent labeling and the red clone-specific knockout of Sox2. We then lineage traced the wild-type (yellow) and knockout (red) cells and quantified the sizes of their clones over time. If Sox2 was essential for stem cell maintenance, we would have expected that the mutant clones would be rapidly lost from the basal layer within a short period of time. By contrast, if Sox2 was not essential for stem cell maintenance, we would have expected that the mutant clones would remain present in the basal layer — and might only be lost due to their relative clonal fitness in that tissue.

**Figure 5.**
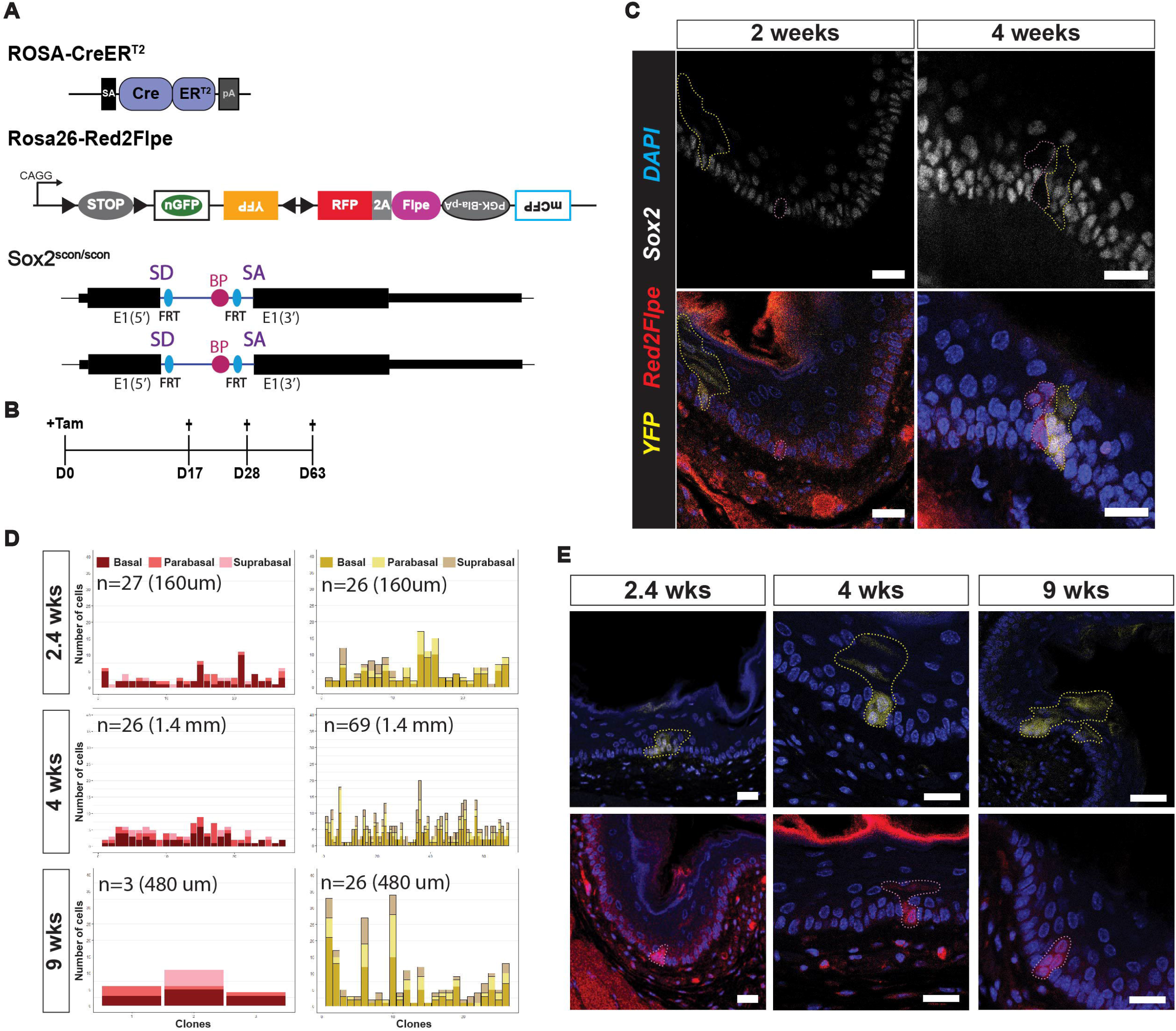
Mosaic knockout of Sox2 reveals its role in the adult esophagus. **A.** Experimental set up for mosaic *Sox2* knockout in an adult mouse esophagus using the *Rosa-CreER^T2^; Red2Flpe; Sox2^scon/scon^* strain. **B.** A single dose of 3mg tamoxifen was injected and the esophagus samples were then harvested on days 17, 28 and 63 after the injection. **C.** Sox2 staining in esophageal sections at 2- and 4-weeks post-induction. The RFP and YFP clones are marked by dashed lines. Scale bar, 20 μm. **D.** Quantification of clone size and spatial distribution of RFP and YFP clones at 2.4-, 4- and 9-weeks post-induction. **E.** Representative images of RFP and YFP clones at 2.4-, 4- and 9-weeks post-induction. The RFP and YFP clones are marked by dashed lines. Scale bar, 20 μm.

We collected the esophagus samples at different time points following the tamoxifen injection (Figure 5B). We first confirmed that Sox2 expression was absent from the RFP+ clones in the esophagus at two weeks and four weeks, and checked that the YFP+ clones expressed Sox2, as expected (Figure 5C). Interestingly, the Sox2 knockout mutant clones remained in the tissue, even after long-term tracing, which suggested that Sox2 might not be essential for stem cell maintenance in the esophagus (Figure 5D and 5E). At 2.4 weeks after induction, the RFP+ and YFP+ clones were found at similar levels (Figure 5D). However, at 4 and 9 weeks, the number of RFP clones was reduced compared with the wild-type YFP+ clones. We also quantified the clone sizes, and the location of the labeled cells within each clone — noting whether they were in the basal, parabasal (stratifying) or suprabasal layers. We found that the size of the RFP+ clones was consistently smaller than the size of the YFP+ clones (Figure 5D). In addition, the RFP+ clones mainly consisted of cells in the basal and parabasal layers, whereas the cells in the YFP+ clones were more evenly distributed across all three layers (Figure 5D and 5E). This suggests that, while the wild-type cells were able to efficiently self-renew and differentiate, the Sox2 mutant cells had reduced capacity for proliferation and differentiation. This eventually led to the loss of RFP+ clones from constant niche competition due to a reduction in their fitness compared to the wild-type cells.

We also utilized the esophageal organoid system to examine the behavior of Sox2 mutant clones over time. We observed a reduced proliferation rate of the Sox2 mutant cells. We also found that the organoid-forming efficiency was lower in Sox2 mutant cells compared to wild-type cells, with smaller organoid sizes observed for all three culture medium conditions tested (WENR+Nic, ENR, or EN) (Figure S5). Nonetheless, Sox2 mutant organoids could still be passaged with both WENR+Nic and ENR media, suggesting persistent stem cell activity. Consistent with the in vivo lineage tracing results above, the esophageal cells that were depleted for Sox2 still retained their stem cell characteristics but exhibited reduced capacity for proliferation.

Sox2 is expressed in the developing foregut and remains expressed in the epithelium of the stomach as well as the esophagus. Therefore, we also checked whether knockout of Sox2 would have an impact on clonal fitness in the stomach epithelium. We found that both the wild-type (YFP+) and the Sox2-mutant (RFP+) glands showed comparable labeling four weeks after being induced (Figure S6A). We also observed the comparable presence of both wild-type and Sox2-mutant clones in the base (Figure S6B and S6C) and isthmus (Figure S6D and S6E) parts of the glands, as we reported previously (Han et al. 2019). Overall, this indicated that although Sox2 may affect cellular differentiation, it does not appear to significantly alter stem cell maintenance or clonal fitness in the stomach epithelium.

## Discussion

Mosaic knockouts are preferred for investigating gene function, as they provide a physiologically normal background for mutant cells; whereas ordinary conditional knockout approaches are associated with risks of strong secondary and tertiary effects, which can obscure the function of the target gene. Intriguingly, when we knocked out Sox2 in a mosaic manner in adult mice we found that Sox2 is not essential for long-term clonal maintenance in the epithelium of the esophagus and stomach. Sox2 is widely known as a stem cell transcription factor in ESCs; however, in the adult esophagus, Sox2 has a more specific role: regulating the proliferation and differentiation of basal cells. Based on the quantification of Sox2 mutant cell behaviors, we found that loss of Sox2 did not abrogate stemness completely, and clonal loss was mainly due to the reduced fitness during cell-cell competition. This observation was confirmed by in vitro organoid culture, where Sox2-mutant esophageal organoids could still be maintained, despite a reduced capacity for proliferation. Our mosaic analyses provide insight into novel aspects of biology that have previously been inaccessible.

Despite significant technical improvements, the generation of cKO mouse lines by zygote injection remains challenging due to the requirement for long template insertion. With our SCON method, we generated several cKO lines using cloning-free reagents and a one-step zygote injection for both Cre-loxP and Flp-FRT recombination systems. SCON alleles are functional in vivo and are compatible with the *Red2Flpe* line for mosaic knockout lineage tracing in the mouse.

In summary, we offer Red2Flpe-SCON as an efficient mosaic genetics tool for studying both wild-type and mutant cell behavior in the same tissue, while minimizing secondary effects. As this system is based on the widely used *Confetti* allele, versatile, tunable, precise, Flp-based mosaic knockouts with multicolor labeling can now be easily achieved in mice (Table 1).

## Materials and methods

### Mice

All animal experiments were performed according to the guidelines of the Austrian Animal Experiments Act; with valid project licenses approved by the Austrian Federal Ministry of Education, Science and Research; and monitored by the institutional IMBA Ethics and Biosafety department.

In this study, Red2Flpe and Apc-FRT mice were generated via embryonic stem cell (ESC) targeting and blastocyst injection. The Red2Flpe mouse was made by targeting in Confetti ESCs of C57BL6/129F1 background, and the Apc-FRT mouse with exon 15 flanked by FRT sites was generated by targeting in ESCs of B6129SF1/J background. Clonal-derived ESCs were injected into C57BL6 blastocysts. Chimeras were then backcrossed to C57BL6 to confirm germline transmission.

### Administering Tamoxifen and harvesting organs for tissue imaging

A dose of either Tamoxifen (Sigma, T5648) dissolved in corn oil (Sigma, C8267), or corn oil only, was injected intraperitoneally into 8–12-week-old mice, with a final concentration of 3 mg tamoxifen per 20 g of body weight. Tissues were harvested, cleaned with cold PBS, and then fixed with 4% paraformaldehyde overnight (16 hours) at 4□. The samples were then washed with cold PBS with 2-3 hours intervals. Intestine samples were embedded in 4% low melting point agarose (Sigma, A9414) and then sectioned using a LAICA VT 1000S vibratome to produce 150 μm thick sections. Esophagus, stomach, spleen, tongue and seminal vesicle samples were incubated in 30% sucrose overnight at 4□, and then frozen on dry ice in O.C.T. solution (Scigen, 4586). The frozen section blocks were cryo-sectioned at 140 μm and then placed in PBS for further processing.

### Immunohistochemistry

Staining was carried out according to methods previously described by Yum et al. (2021)Yum et al. (2021)Yum et al. (2021). Briefly, samples were incubated in a blocking solution (5%DMSO, 0.5% Triton X-100, and 2% normal donkey serum (NDS) in PBS) for 4 hours at *4¤* on a shaker. The solution was then exchanged with primary antibody diluted in a staining solution (1% DMSO, 0.5% Triton X-100, and 2% NDS in PBS): Sox2 (1:200; Invitrogen, 14-9811-82) or AF647-conjugated β-catenin (1:200; Cell Signaling Technology, 4627S). Sections were then incubated for 48 hours at *4□* on a shaker. After incubation, the samples were washed three times in cold PBS with 2-3 hour intervals between each wash. The samples were then stained with secondary antibodies (1:500; donkey anti-rat AF647, Invitrogen) diluted in staining solution and incubated for 48 hours at *4□* on a shaker. The samples were then washed in PBS three times and stained with DAPI overnight at 4□. After washing, the samples were transferred to microscopic slides and mounted in RapiClear1.52 (Sunjin Lab). All confocal images were taken using a multiphoton SP8 confocal microscope (Leica).

### Preparation of primary intestinal cells for fluorescence-activated cell sorting (FACS)

The proximal halves of the freshly isolated intestines were gently cleaned by flushing them through cold PBS. The tissues were opened longitudinally and then shaken vigorously in cold PBS to clean the samples further. Once the solution became clear, the samples were incubated in Gentle Cell Dissociation Reagent (100-0485, STEMCELL technologies) for 30 minutes on ice. The samples were then shaken vigorously to release the cells from the tissue. The solution was then filtered through a 70 μm filter, and then centrifuged at 300g for 5 minutes at 4°C. Following this, the cell pellet was resuspended in TrypLE Express Enzyme (12604013, Gibco^TM^) and incubated at 37°C for 5 minutes to dissociate the cell clumps into single cells. Cold PBS was then added to the single cell suspension. The cell suspension was then centrifuged at 300g for 5 minutes at 4°C. Following this, the resulting cell pellet was resuspended in FACS buffer (2% FBS, 2mM EDTA in PBS) and filtered using 40 μm strainers. The cells were analyzed using a BD-LSRFortessa flow cytometer (BD), and the flow cytometry data were analyzed using FlowJo software (BD).

### eG-SCON-FRT-FP constructs

The eG-SCON-FRT-FP cassette was ordered from Genscript, and subsequently cloned into pcDNA4/TO, and a piggybac overexpressing the plasmid. The pcDNA4/TO-eG-SCON-FRT-FP vectors were then transformed into Flp-expressing bacteria (A710, Gene bridges) to obtain the recombined forms. The correct clones were confirmed by restriction digest and subsequent Sanger sequencing.

### Cell cultures and transfections

#### HEK293T cells

Human embryonic kidney (HEK) 293T cells were cultured in high glucose DMEM containing 10% fetal bovine serum (FBS, Sigma), 1% penicillin-streptomycin (P/S; Sigma, P0781) and 1% L-glutamine (L-glut; Gibco, 25030024).

#### Mouse ESCs

The mouse ES cell line AN3-12 was cultured and transfected as previously described by Wu et al. (2022), when they tested the recombination compatibility of SCON-FRT. Confetti and Red2Flpe ESCs were cultured in cell culture-grade dishes pre-coated with 10% gelatin (Sigma, G1890), and then supplied with DMEM/F12 (Sigma, D6421), Neurobasal (Gibco, 21103049), N2 (Gibco, 17502048), B27 (Gibco, 17504044), 1% P/S (Sigma, P0781), 0.1 mM 2-mercaptoethanol (Sigma, M7522), 1% L-glut (Gibco, 25030024), 10 mM PD0325901 (Axon, 1408), 10 mM CHIR99021 (Axon, 1386), and 10^3^ units/ ml of mouse ESGRO LIF (Merck Millipore ESG1107).

#### Plasmid transfection to Hek293T cells

Transfection and subsequent analyses were performed as previously described by Wu et al. (2022)Wu et al. (2022)Wu et al. (2022). Briefly, 500,000-750,000 Hek293T cells were seeded in 6-well plates and left to attach and grow overnight. Following this, 2.5 μg of DNA (1 μg of mCherry-expressing plasmid (Addgene, 72264), and 1.5 μg of either pcDNA4/TO-eGFP, -eG-SCON-FRT-FP, or recombined forms of eG-SCON-FRT-FP was mixed with 8 μl of polyethyleneimine (1 mg/ml; Polysciences, 23966). These solutions were then incubated at room temperature for at least 15 minutes before being added dropwise to the cells. The culture medium was exchanged after 8-12 hours or after an overnight transfection. The cells were examined 24–36 hours after transfection, under an EVOS M7000 microscope (Thermo Scientific) using brightfield, GFP and TexasRed filters. Then, 36–48 hours after transfection, cells were dissociated into single cells for flow cytometry analysis, with a BD-LSRFortessa flow cytometer (BD). Data from the flow cytometry experiments were analyzed using FlowJo software (BD).

### Organoid cultures

#### Establishment and maintenance of esophageal organoids

The esophagus harvested from an 8–12-week-old mouse was cut open longitudinally and incubated in 50 mM EDTA for 5 minutes at 37□. The epithelial layer was carefully peeled from the stroma, minced into small pieces and then incubated for 10 minutes at 37□ in 0.5 mg/ml of Dispase dissolved in DMEM (Sigma; D4818). The solution containing the esophageal pieces was diluted in PBS and pipetted vigorously using a P1000 pipette before being filtered using a 30 μm cell strainer. The cells were spun down at 300g for 5 minutes and then resuspended in cold PBS. This step was then repeated before embedding the cells in basement membrane extract (BME-R1) (R&D Systems, 3433010R1). Droplets measuring 15 μl were placed in a 48-well plate (Sigma, CLS3548-100EA) and transferred to a 37□ incubator for 3-5 minutes until the BME had polymerized.

Cells were supplemented with WENR+Nic medium, which consisted of advanced DMEM/F12 (Gibco, 12634028) supplemented with P/S (1%; Sigma, P0781), 10 mM HEPES (Gibco, 15630056), Glutamax (1%; Gibco, 35050061), B27 (2%; Life Technologies, 17504044), Wnt3 conditioned medium (Wnt3a L-cells, 50% of final volume), 50 ng/ml recombinant mouse epidermal growth factor (EGF; Gibco, PMG8041), 100 ng/ml recombinant murine Noggin (PeproTech, 250-38), R-spondin-1 conditioned medium (HA-R-Spondin1-Fc 293T cells, 10% of final volume), and 10 mM nicotinamide (MilliporeSigma, N0636). This medium was used for the first two passages and was then changed to ENR media. Organoids were passaged weekly by collecting BME droplets in DMEM/F12 and pipetting several times to break up the droplets. They were then spun down at 900g for 3 minutes. The supernatant was discarded carefully using a pipette, and the organoids were then resuspended in TrypLE Express (Gibco). They were then incubated at 37 □ for 10-15 minutes. Following this, the solution was pipetted using a p200 pipette to facilitate dissociation. The cells were then examined under the microscope. An aliquot of the cell solution was then transferred to a new tube to obtain a 1:10 split ratio. This was spun down at 900g for 3 minutes, and then embedded in BME.

#### Treatment with *4-Hydroxytamoxifen*

After two or more passages, esophageal organoids, cultured from single cells in the presence of EGF, Noggin, and R-Spondin 1 (collectively known as ENR), were grown for four days to allow organoids to expand in size. Medium containing vehicle (ethanol) or 500 nM 4-OH-tamoxifen (Sigma, H7904) was added after BME polymerized. After 8 hours, the medium was changed back to ENR, and then replenished every two days. One week after the 4-OH Tam induction, the organoids were dissociated into single cells using TrypLE Express (Gibco). Cells were then sorted using a Sony SH800 cell sorter, equipped with 405nm, 488nm and 561nm lasers (Sony).

#### Imaging

The cultured organoids were examined under an EVOS M7000 microscope (Thermo Scientific) using brightfield, GFP and RFP filters.

## Acknowledgements

We thank present and past members of the Koo, Elling and Urban labs at IMBA for valuable discussions and critical comments, Dr. Rike Zietlow and the Life Science Editors for reading and correcting the manuscript, VBC core facilities (especially the Histopathology facility, BioOptics and the animal caretakers). This work was supported by core funding from the Institute of Molecular Biotechnology (IMBA) of the Austrian Academy of Sciences; ERC starting grant, Troy Stem cells, 639050; Interpark Bio-Convergence Center Grant Program; and fellowship to S.W. (DOC Fellowship of the Austrian Academy of Sciences) and G.C. (Lise Meitner Postdoctoral fellowship M2976, FWF).

## Author contributions

B.-K.K, Joo-Hyeon L. and B.D.S. formulated the design of Red2cDNA. S.W. and B.-K.K. planned and designed the experiments. R.B. and S.W. performed SCON-FRT transfection experiments and the subsequent FACS analysis. Ji-Hyun L. and S.W. analyzed the stomach phenotype of mosaic Sox2 knockout. S.W. performed all experiments, with help from G.C., S.Y.P., I.T., J.K., N.H. and S.P.C, and the core facilities at IMBA/IMP. C.T. performed blastocyst injection of targeted mouse ESCs to generate Red2Flpe and ApcFRT mice. S.K., H.L., and J.K.K. performed all computational analysis.

---

**Figure S1.**
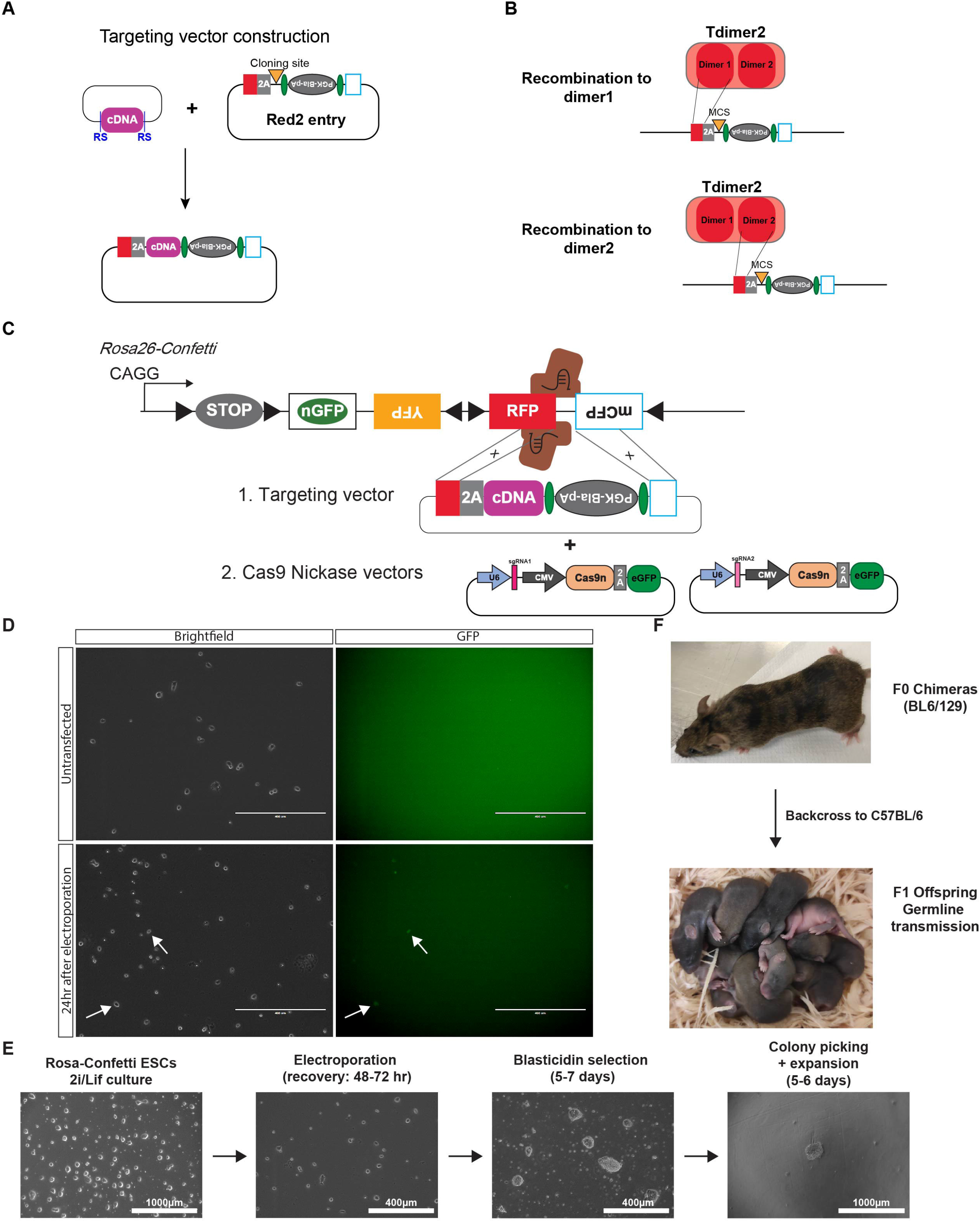
Pipeline for generating new Red2cDNA mouse lines. **A.** The Red2 targeting vector consists of homology arms that correspond to parts of the RFP and mCFP sequences, with a PGK-blasticidin-pA cassette in a reversed orientation, flanked by FRT sites (in green), and a cloning site for any cDNA sequence downstream of a P2A peptide. **B.** The left homology arm of the Red2 targeting vector can be recombined to either one of the RFP (tdimer2) dimer sequences. **C.** Targeting onto the confetti allele is achieved with a pair of Cas9 nickase vectors, that contain gRNA targeted to RFP at the 3’ terminus. **D.** Successful delivery of the targeting components can be checked 24hrs post-electroporation, where eGFP expression from the Cas9 nickase vectors can be observed in a small subset of cells. **E.** Steps for ESC targeting (which requires 2.5-3 weeks), including initial expansion of ESCs, electroporation, blasticidin selection, colony picking, and expansion. **F.** Chimeric mice are generated after the correct targeting clones (B6/129F1 background) have been injected into a developing blastocyst (C57BL/6), and back crossing to confirm germline transmission (agouti pups) has been completed.

**Figure S2.**
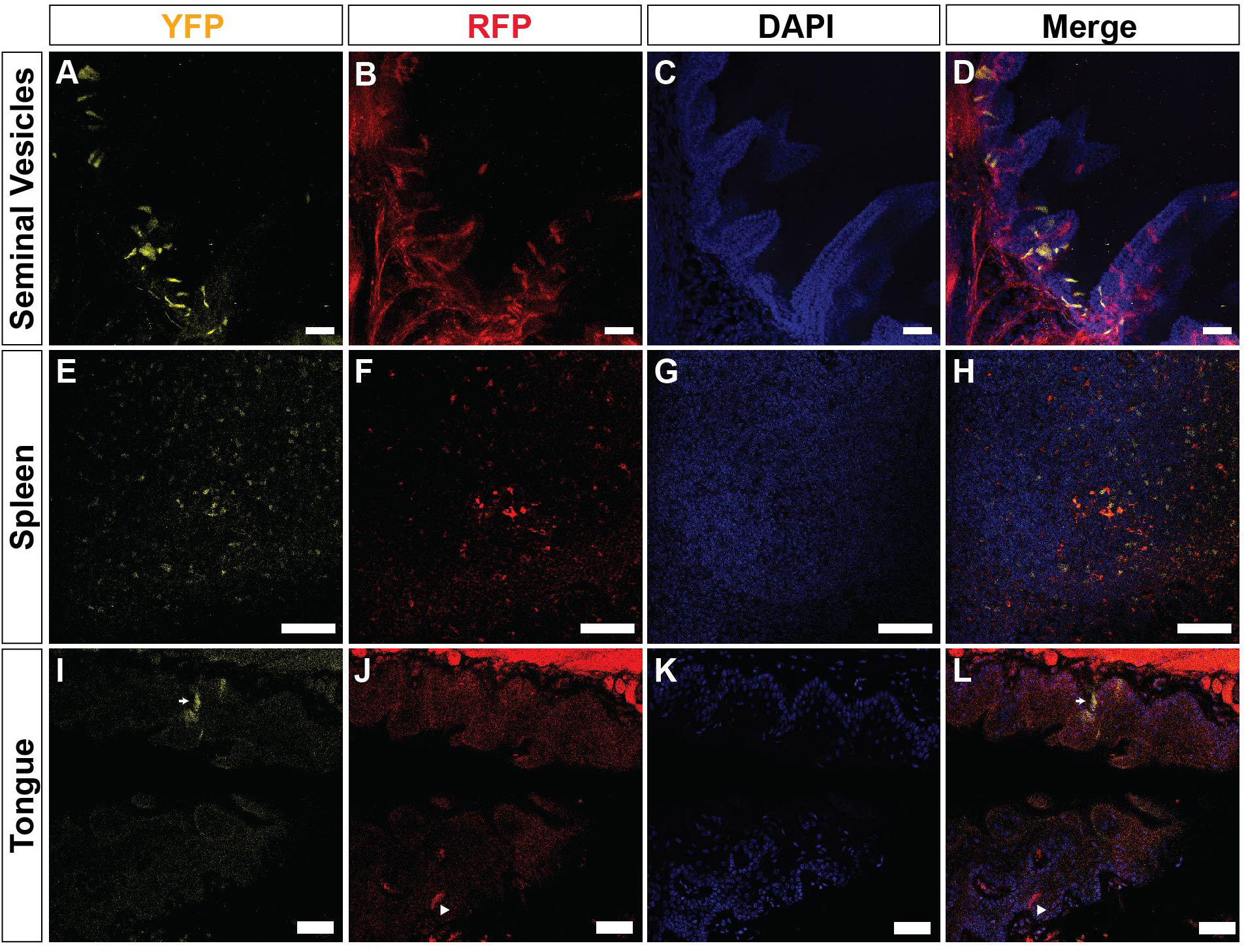
Usability of Red2Flpe in multiple organs. **A-D.** Seminal vesicles. Scale bar, 30 μm. **E-H,** Spleen. Scale bar, 30 μm. **I-L,** Tongue. Scale bar, 20 μm. RFP and YFP cells are marked by triangles and arrows, respectively.

**Figure S3.**
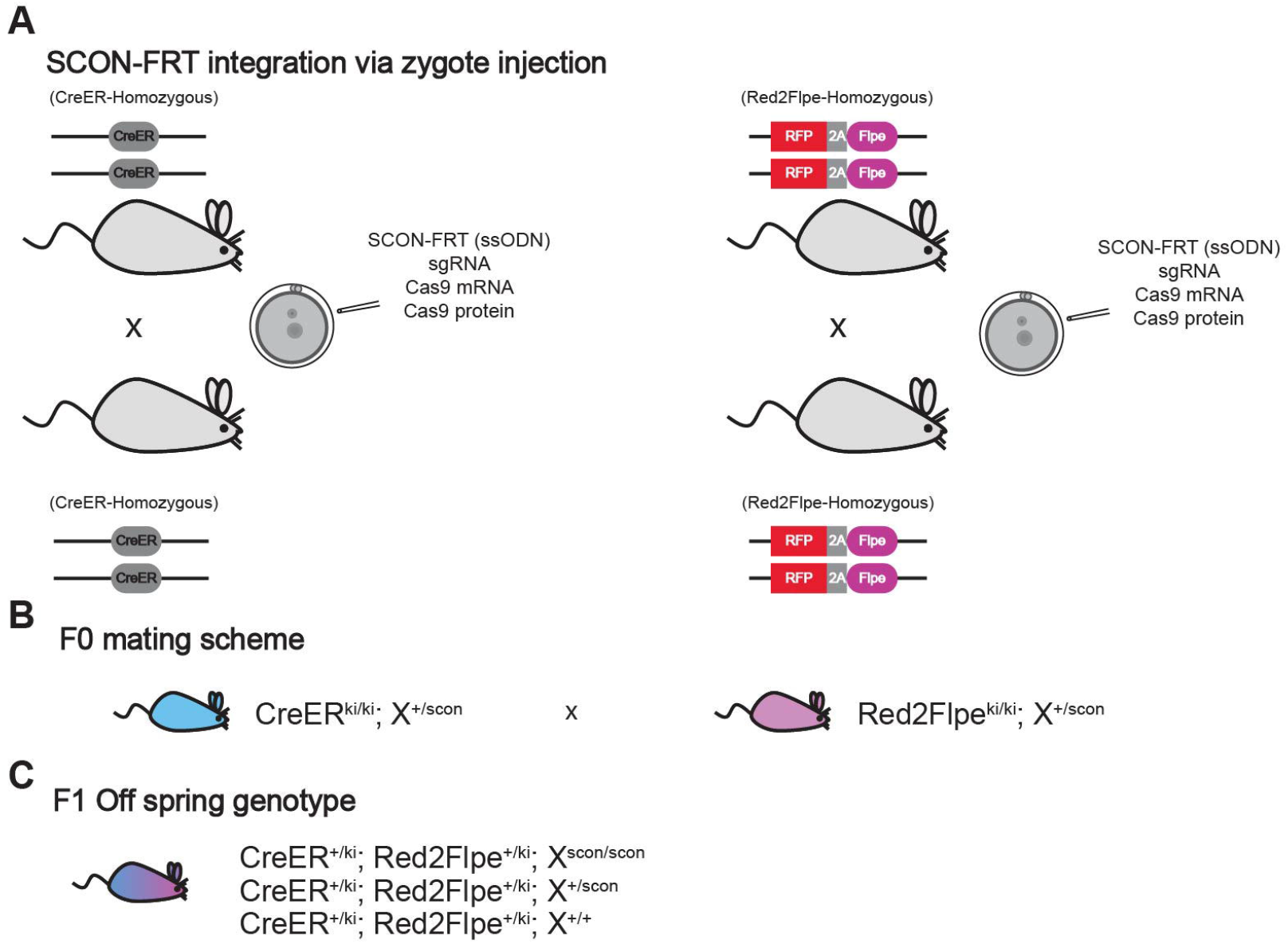
Generation of mosaic conditional mice via zygote injection. **A.** Parallel zygote targeting of SCON-FRT to target the gene in the desired CreER and Red2Flpe homozygous mice. **B.** All offspring contain homozygous CreER and Red2Flpe. Pups containing precise SCON-FRT integration in the gene of interest (gene X) are bred to obtain experimental cohorts and confirm germline transmission of X^scon^. **C.** There is a ¼ chance of obtaining experimental cohorts containing CreER (heterozygous), Red2Flpe (heterozygous) and SCON-FRT (homozygous).

**Figure S4.**
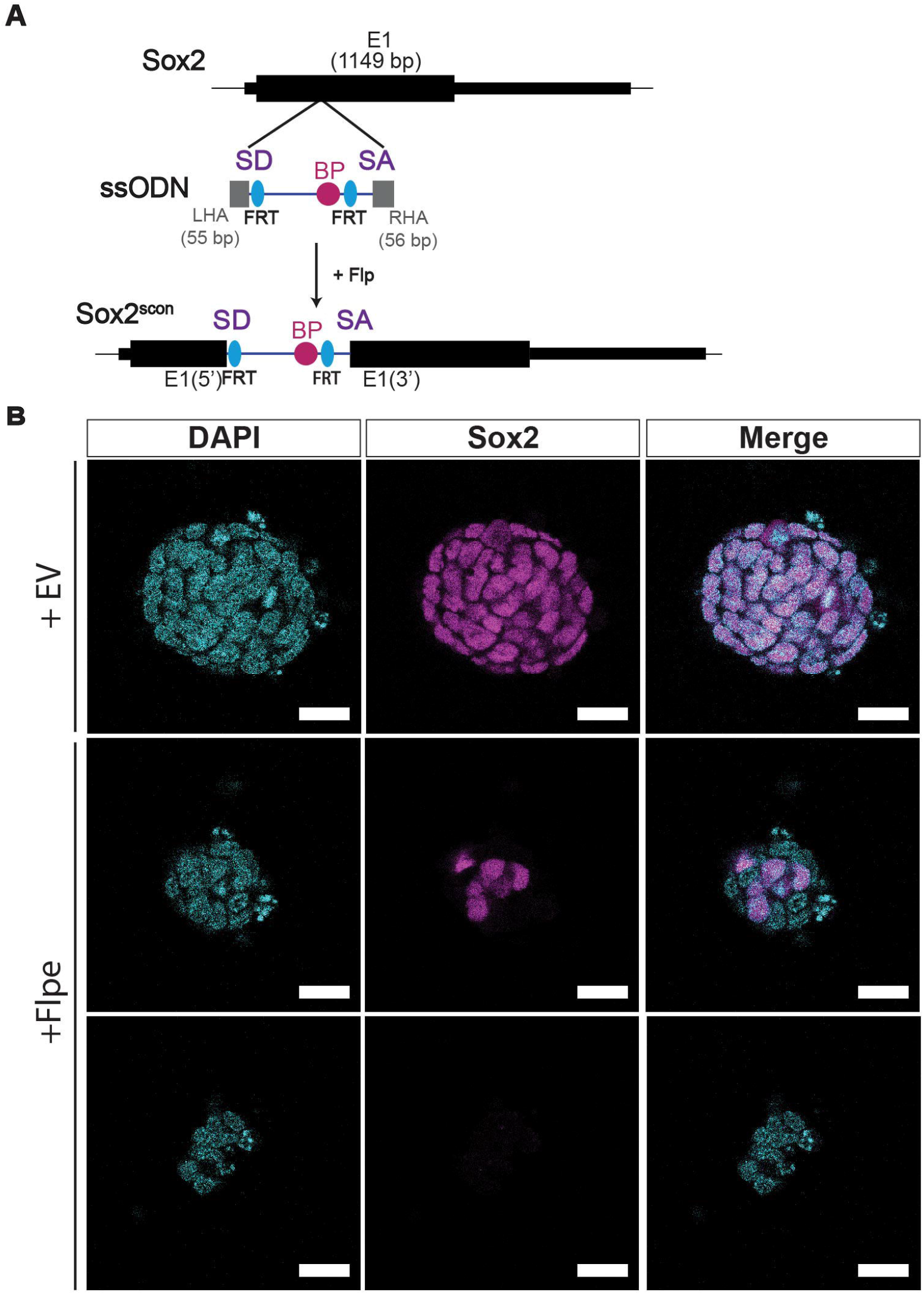
Sox2-SCONFRT: A new cKO mouse line that functions as expected. **A.** Design of the Sox2-SCON-FRT allele. The synthesized ssODN of the 189bp long SCON-FRT cassette, with homology arms of 55–56 bp, is inserted into the Sox2 allele via zygote injection. **B.** Images of Sox2-SCON-FRT homozygous mouse ESCs 48 hours after transfection with either the Flpe-expressing plasmid or an empty vector control. Scale bar, 20 μm.

**Figure S5.**
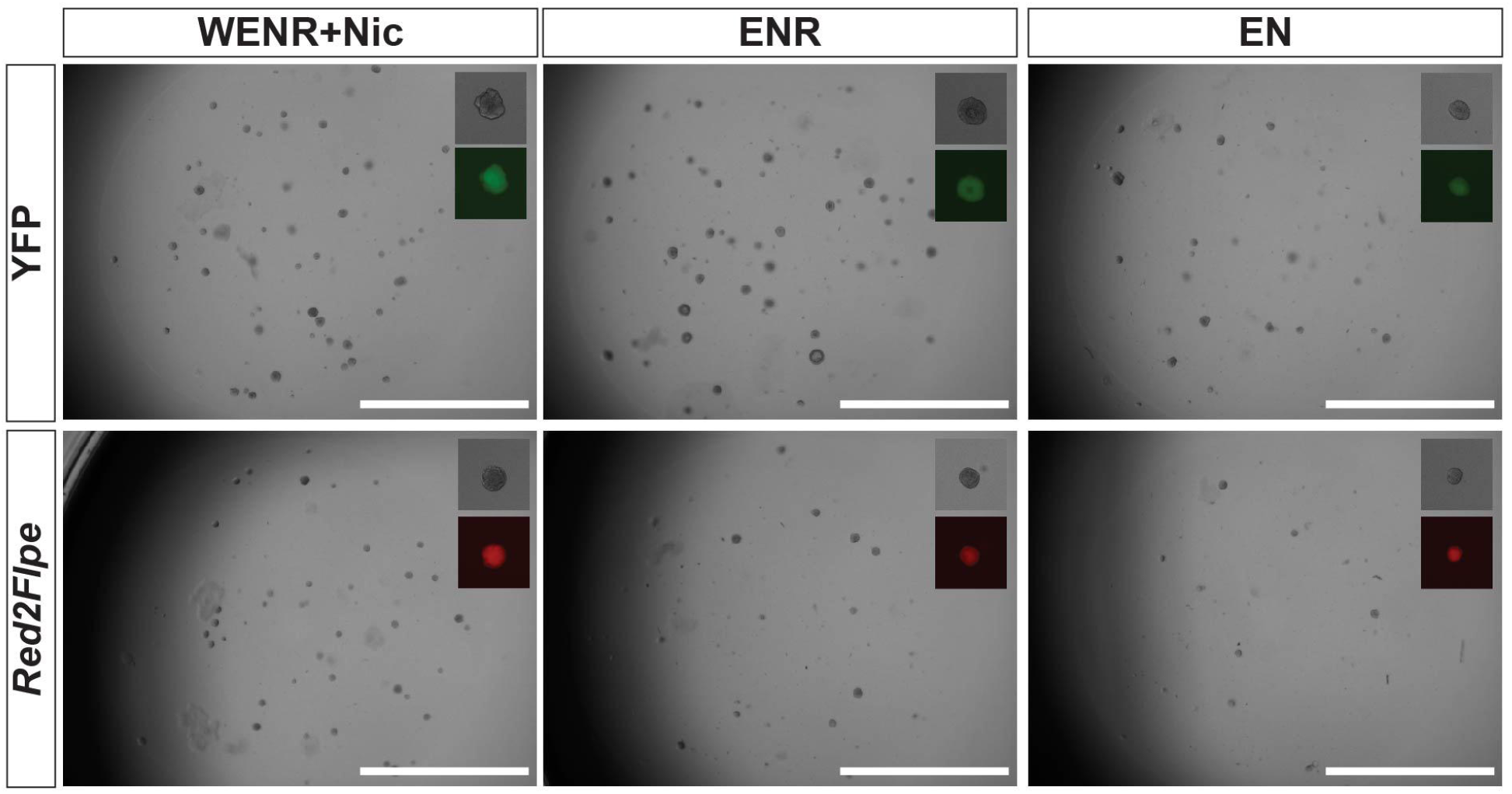
Sox2 knockout esophageal cells show reduced self-renewal and organoid forming efficiency. **A.** RFP+ and YFP esophageal organoids of Rosa-CreER^T2(+)^, Red2Flpe^+^, and Sox2^sc/sc^, cultured in WENR+Nic (Wnt3a-condition medium, EGF, Noggin, R-spondin1 and nicotinamide), ENR (EGF, Noggin and R-spondin1) or EN (EGF and Noggin). Scale bar, 2 mm.

**Figure S6.**
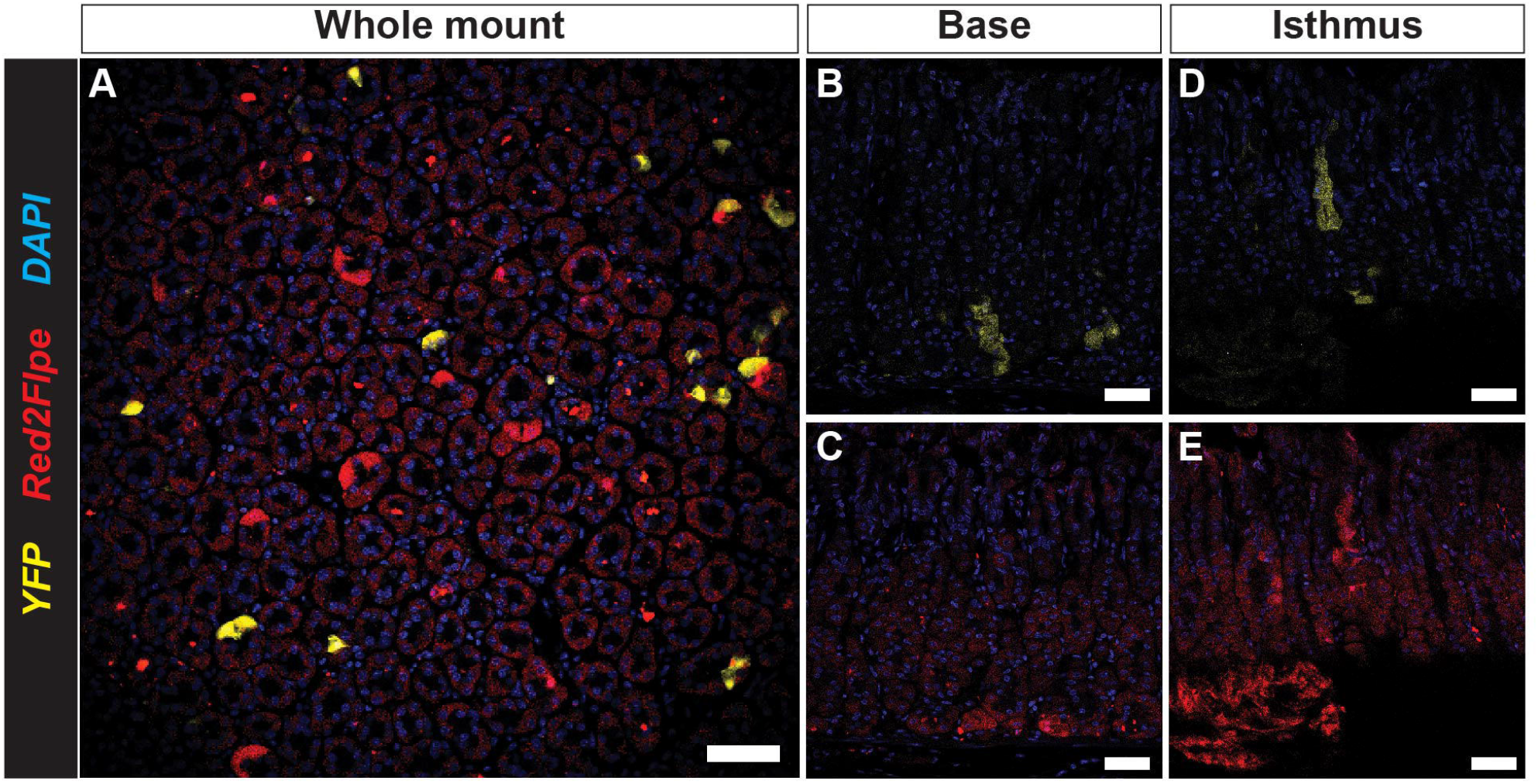
Sox2 mosaic knockout in the adult stomach 4 weeks after induction. **a.** Wholemount of the corpus epithelium reveals clonality progression of the glands. Scale bar, 40 μm. **b-c,** Basal clones of YFP and RFP cells. Scale bar, 20 μm. **d-e**, Isthmus clones of YFP and RFP cells. Scale bar, 20 μm.

## References

Arnold K, Sarkar A, Yram MA, et al (2011) Sox2 + adult stem and progenitor cells are important for tissue regeneration and survival of mice. Cell Stem Cell 9:317–329. https://doi.org/10.1016/j.stem.2011.09.001

Buchholz F, Angrand P, Stewart AF (1998) Evolved By Cycling N1Utagenesis. Nat Biotechnol 16:657–662

Campbell RE, Tour O, Palmer AE, et al (2002) A monomeric red fluorescent protein. Proceedings of the National Academy of Sciences 99:7877–7882. https://doi.org/10.1073/pnas.082243699

Colnot S, Niwa-Kawakita M, Hamard G, et al (2004) Colorectal cancers in a new mouse model of familial adenomatous polyposis: influence of genetic and environmental modifiers. Laboratory Investigation 84:1619–1630. https://doi.org/10.1038/labinvest.3700180

Colom B, Herms A, Hall MWJ, et al (2021) Mutant clones in normal epithelium outcompete and eliminate emerging tumours. Nature 598:510–514. https://doi.org/10.1038/s41586-021-03965-7

Contreras X, Amberg N, Davaatseren A, et al (2021) A genome-wide library of MADM mice for single-cell genetic mosaic analysis. Cell Rep 35:. https://doi.org/10.1016/j.celrep.2021.109274

DeWard AD, Cramer J, Lagasse E (2014) Cellular Heterogeneity in the Mouse Esophagus Implicates the Presence of a Nonquiescent Epithelial Stem Cell Population. Cell Rep 9:701–711. https://doi.org/10.1016/j.celrep.2014.09.027

Doupé DP, Alcolea MP, Roshan A, et al (2012) A Single Progenitor Population Switches Behavior to Maintain and Repair Esophageal Epithelium. Science (1979) 337:1091–1093. https://doi.org/10.1126/science.1218835

Han S, Fink J, Jörg DJ, et al (2019) Defining the Identity and Dynamics of Adult Gastric Isthmus Stem Cells. Cell Stem Cell 25:342–356.e7. https://doi.org/10.1016/j.stem.2019.07.008

Kim GB, Rincon Fernandez Pacheco D, Saxon D, et al (2019) Rapid Generation of Somatic Mouse Mosaics with Locus-Specific, Stably Integrated Transgenic Elements. Cell 179:251–267.e24. https://doi.org/10.1016/j.cell.2019.08.013

Kohara K, Inoue A, Nakano Y, et al (2020) BATTLE: Genetically Engineered Strategies for Split-Tunable Allocation of Multiple Transgenes in the Nervous System. iScience 23:101248. https://doi.org/10.1016/j.isci.2020.101248

Lao Z, Raju GP, Bai CB, Joyner AL (2012) MASTR: A Technique for Mosaic Mutant Analysis with Spatial and Temporal Control of Recombination Using Conditional Floxed Alleles in Mice. Cell Rep 2:386–396. https://doi.org/10.1016/j.celrep.2012.07.004

Lee-Six H, Øbro NF, Shepherd MS, et al (2018) Population dynamics of normal human blood inferred from somatic mutations. Nature 561:473–478. https://doi.org/10.1038/s41586-018-0497-0

Lopez-Garcia C, Klein AM, Simons BD, Winton DJ (2010) Intestinal Stem Cell Replacement Follows a Pattern of Neutral Drift. Science (1979) 330:822–825. https://doi.org/10.1126/science.1196236

Martincorena I, Fowler JC, Wabik A, et al (2018) Somatic mutant clones colonize the human esophagus with age. Science (1979) 917:911–917. https://doi.org/10.1126/science.aau3879

Martincorena I, Roshan A, Gerstung M, et al (2015) High burden and pervasive positive selection of somatic mutations in normal human skin. Science (1979) 348:880–886. https://doi.org/10.1126/science.aaa6806

Masui S, Nakatake Y, Toyooka Y, et al (2007) Pluripotency governed by Sox2 via regulation of Oct3/4 expression in mouse embryonic stem cells. Nat Cell Biol 9:625–635. https://doi.org/10.1038/ncb1589

Pontes-Quero S, Heredia L, Casquero-García V, et al (2017) Dual ifgMosaic: A Versatile Method for Multispectral and Combinatorial Mosaic Gene-Function Analysis. Cell 170:800–814.e18. https://doi.org/10.1016/j.cell.2017.07.031

Quadros RM, Miura H, Harms DW, et al (2017) Easi-CRISPR: A robust method for one-step generation of mice carrying conditional and insertion alleles using long ssDNA donors and CRISPR ribonucleoproteins. Genome Biol 18:1–15. https://doi.org/10.1186/s13059-017-1220-4

Que J, Okubo T, Goldenring JR, et al (2007) Multiple dose-dependent roles for Sox2 in the patterning and differentiation of anterior foregut endoderm. Development 134:2521–2531. https://doi.org/10.1242/dev.003855

Ran FA, Hsu PD, Lin C-Y, et al (2013) Double Nicking by RNA-Guided CRISPR Cas9 for Enhanced Genome Editing Specificity. Cell 154:1380–1389. https://doi.org/10.1016/j.cell.2013.08.021

Sarkar A, Hochedlinger K (2013) The Sox family of transcription factors: Versatile regulators of stem and progenitor cell fate. Cell Stem Cell 12:15–30. https://doi.org/10.1016/j.stem.2012.12.007

Snippert HJ, Schepers AG, Van Es JH, et al (2014) Biased competition between Lgr5 intestinal stem cells driven by oncogenic mutation induces clonal expansion. EMBO Rep 15:62–69. https://doi.org/10.1002/embr.201337799

Snippert HJ, van der Flier LG, Sato T, et al (2010) Intestinal crypt homeostasis results from neutral competition between symmetrically dividing Lgr5 stem cells. Cell 143:134–144. https://doi.org/10.1016/j.cell.2010.09.016

Sousa VH, Miyoshi G, Hjerling-Leffler J, et al (2009) Characterization of Nkx6-2-derived neocortical interneuron lineages. Cerebral Cortex 19:1–10. https://doi.org/10.1093/cercor/bhp038

Thorsen AS, Khamis D, Kemp R, et al (2021) Heterogeneity in clone dynamics within and adjacent to intestinal tumours identified by Dre-mediated lineage tracing. DMM Disease Models and Mechanisms 14:. https://doi.org/10.1242/DMM.046706

Trisno SL, Philo KED, McCracken KW, et al (2018) Esophageal Organoids from Human Pluripotent Stem Cells Delineate Sox2 Functions during Esophageal Specification. Cell Stem Cell 23:501–515.e7. https://doi.org/10.1016/j.stem.2018.08.008

Vermeulen L, Morrissey E, Van Der Heijden M, et al (2013) Defining stem cell dynamics in models of intestinal tumor initiation. Science (1979) 342:995–998. https://doi.org/10.1126/science.1243148

Wu SH, Lee JH, Koo BK (2019) Lineage tracing: Computational reconstruction goes beyond the limit of imaging. Mol Cells 42:104–112

Wu S-HS, Lee H, Szép-Bakonyi R, et al (2022) SCON—a Short Conditional intrON for conditional knockout with one-step zygote injection. Exp Mol Med. https://doi.org/10.1038/s12276-022-00891-0

Yum MK, Han S, Fink J, et al (2021) Tracing oncogene-driven remodelling of the intestinal stem cell niche. Nature 594:442–447. https://doi.org/10.1038/s41586-021-03605-0

Zong H, Espinosa JS, Su HH, et al (2005) Mosaic analysis with double markers in mice. Cell 121:479–492. https://doi.org/10.1016/j.cell.2005.02.012

